# Biophysical carbon concentrating mechanisms in land plants: insights from reaction-diffusion modeling

**DOI:** 10.1101/2024.01.04.574220

**Authors:** Joshua A.M. Kaste, Berkley J. Walker, Yair Shachar-Hill

## Abstract

Carbon Concentrating Mechanisms (CCMs) have evolved numerous times in photosynthetic organisms. They elevate the concentration of CO_2_ around the carbon-fixing enzyme rubisco, thereby increasing CO_2_ assimilatory flux and reducing photorespiration. Biophysical CCMs, like the pyrenoid-based CCM of *Chlamydomonas reinhardtii* or carboxysome systems of cyanobacteria, are common in aquatic photosynthetic microbes, but in land plants appear only among the hornworts. To predict the likely efficiency of biophysical CCMs in C3 plants, we used spatially resolved reaction-diffusion models to predict rubisco saturation and light use efficiency. We find that the energy efficiency of adding individual CCM components to a C3 land plant is highly dependent on the permeability of lipid membranes to CO_2_, with values in the range reported in the literature that are higher than used in previous modeling studies resulting in low light use efficiency. Adding a complete pyrenoid-based CCM into the leaf cells of a C3 land plant is predicted to boost net CO_2_ fixation, but at higher energetic costs than those incurred by photorespiratory losses without a CCM. Two notable exceptions are when substomatal CO_2_ levels are as low as those found in land plants that already employ biochemical CCMs and when gas exchange is limited such as with hornworts, making the use of a biophysical CCM necessary to achieve net positive CO_2_ fixation under atmospheric CO_2_ levels. This provides an explanation for the uniqueness of hornworts’ CCM among land plants and evolution of pyrenoids multiple times.

## Introduction

Ribulose-1,5-bisphosphate carboxylase/oxygenase (rubisco) catalyzes the fixation of CO_2_ as part of the Calvin-Benson Cycle (CBC) but is also capable of fixing O_2_. The fixation of O_2_ results in the formation of 2-phosphoglycolate (2PG), with the photorespiratory pathway being necessary to detoxify and recover the carbon in 2PG and recycle it back into the CBC. Although rubisco shows selectivity for CO_2_ relative to O_2_, significant photorespiratory flux still occurs in photosynthetic systems due to the much higher partial pressure of O_2_ in the earth’s atmosphere relative to CO_2_. Photorespiratory flux lowers net carbon assimilation and incurs substantial energetic costs, in the form of ATP, redox equivalents, and ultimately photons. Although the costs associated with photorespiration vary between plant species and environmental conditions, it has been estimated that photorespiration accounts for crop yield decreases of 20 and 36% for soybean and wheat respectively under current climate conditions (Walker et al., 2016).

Carbon Concentrating Mechanisms (CCMs) increase the concentration of CO_2_ around rubisco, competitively inhibiting the oxygenation reaction, suppressing photorespiration, and increasing carboxylation flux (Raven et al., 2017). In biochemical CCMs, such as C4 and CAM photosynthesis, inorganic carbon is fixed into an intermediate form of organic carbon, before eventually being released around rubisco (Ludwig, 2013; Bräutigam et al., 2017). Biophysical or “inorganic” CCMs, on the other hand, do not rely on any additional intermediate organic carbon species, but instead use transport-driven pumps, diffusional barriers, carbonic anhydrases, and pH differences between cellular compartments to increase the CO_2_ concentration near rubisco (Raven et al., 2008). Such CCMs are common in cyanobacteria and algae (Raven et al., 2008), but are conspicuously absent in C3 plants, including almost all land plants. This has motivated researchers to look into the possibility of introducing a CCM, either in its entirety or individual components, into these plants to improve carbon fixation, reduce photorespiratory CO_2_ and energy losses, and ultimately boost yields (Ermakova et al., 2020; Hennacy and Jonikas, 2020).

The seemingly substantial benefits of CCMs raise the question of why they are not already more widespread in land plants. Despite their lack of a CCM, C3 plants are still the most abundant group of land plants in terms of vegetation coverage and gross photosynthetic productivity (Still et al., 2003; Raven et al., 2017). In the case of C4 photosynthesis, the large number of anatomical and biochemical features required has been invoked as a reason why, rather than being universally adopted in land plants, it has instead repeatedly evolved only in lineages exposed to the kinds of hot, arid conditions that limit water availability and exacerbate the losses associated with photorespiration (Sage et al., 2018).

However, such an explanation is less satisfactory in the case of biophysical CCMs because they are present in the hornworts. It also raises the question of why biophysical CCMs are uniformly absent in all land plant lineages except for the hornworts (Villarreal and Renner, 2012).

Have inefficiencies associated with biophysical CCMs precluded their successful emergence in C3 plants and can we examine the presence and absence of these biophysical CCMs in different groups of organisms using these inefficiencies? The efficiency of intermediate photosynthetic configurations, featuring some but not all of the essential parts of a CCM, may also represent a barrier to the emergence of CCMs in land plant lineages. Anatomical and life history details of hornworts may explain why, among the land plants, only hornworts have evolved pyrenoid-based biophysical CCMs (PCCMs), and have done so repeatedly (Villarreal and Renner, 2012). The poikilohydric life history of hornworts makes it necessary for them to have highly desiccation-tolerant cell walls which, together with bryophytes’ generally thicker cell walls (Flexas et al., 2021) and hornworts’ simpler tissue architecture, may explain their extremely low gas conductance (Meyer et al., 2008; Carriquí et al., 2019). We hypothesized that the distinct morphologic characteristics and habitat of hornworts may explain why they, uniquely among the land plants, evolved biophysical CCMs. It is possible that the different paths that inorganic carbon has to take from the environment into a C3 land plant cell versus an algal cell can similarly explain why the former never uses pyrenoids to concentrate carbon and the latter frequently does.

A closer examination of the costs of a CCM may also inform the viability and strategy of biotechnological projects focused on introducing them to C3 crops. Prior quantitative modeling work argues that incorporating individual CCM components – in particular, bicarbonate transporters at the chloroplast membrane – and entire CCMs into land plant systems may boost net CO_2_ fixation as well as improve the efficiency of photosynthetic carbon assimilation by reducing the energetic costs associated with photorespiration (McGrath and Long, 2014; Fei et al., 2022). Similar arguments have been made in favor of engineering biochemical – e.g. C4 – photosynthesis into C3 plants (Walker et al., 2016). These models represent sophisticated, integrative descriptions of photosynthetic carbon assimilation. For the purposes of the questions we are interested in, however, we needed models of both land plant and algal systems that represent photo-assimilatory processes at the whole-cell level. We also needed models that allow us to explore substantial uncertainties in certain key parameters, and that include energy costs associated with carbonic anhydrase (CA) activity in the thylakoid lumen.

Here we developed spatially-resolved reaction-diffusion models of land plants and green algae with and without PCCMs in the *Virtual Cell* platform (Schaff et al., 1997; Cowan et al., 2012). These models represent, to our knowledge, the first such models of C3 land plants containing pyrenoid-based biophysical CCMs, as well as the first models of algal systems containing biophysical CCMs going beyond the scale of the chloroplast and including the whole cell in an aqueous environment. We highlight the substantial uncertainty in reported or predicted values of the permeability of lipid membranes to CO_2_ and explore how this uncertainty can give rise to qualitatively different conclusions as to the efficiency and effectiveness of adding chloroplast envelope bicarbonate pumps in particular. Finally, we find that despite the near-ubiquity of biophysical CCMs in algae, modeling suggests that lower levels of external inorganic carbon (DIC) are needed to make CCMs energetically favorable for land plants.

## Methods

### Model details

Spatially-resolved reaction-diffusion models of carbon assimilation were developed in the *Virtual Cell* platform, a software suite that allows for the creation and analysis of chemical reaction diffusion dynamics in the context of 3D models (Schaff et al., 1997; Cowan et al., 2012). Baseline parameters for simulations can be found in **Table 1** and diagrams of two of the models used in this study, showing the representative features of the land plant and algal models, as well as the differences between the with-and without-PCCM models, can be seen in **Figure 1**.

**Figure 1:**
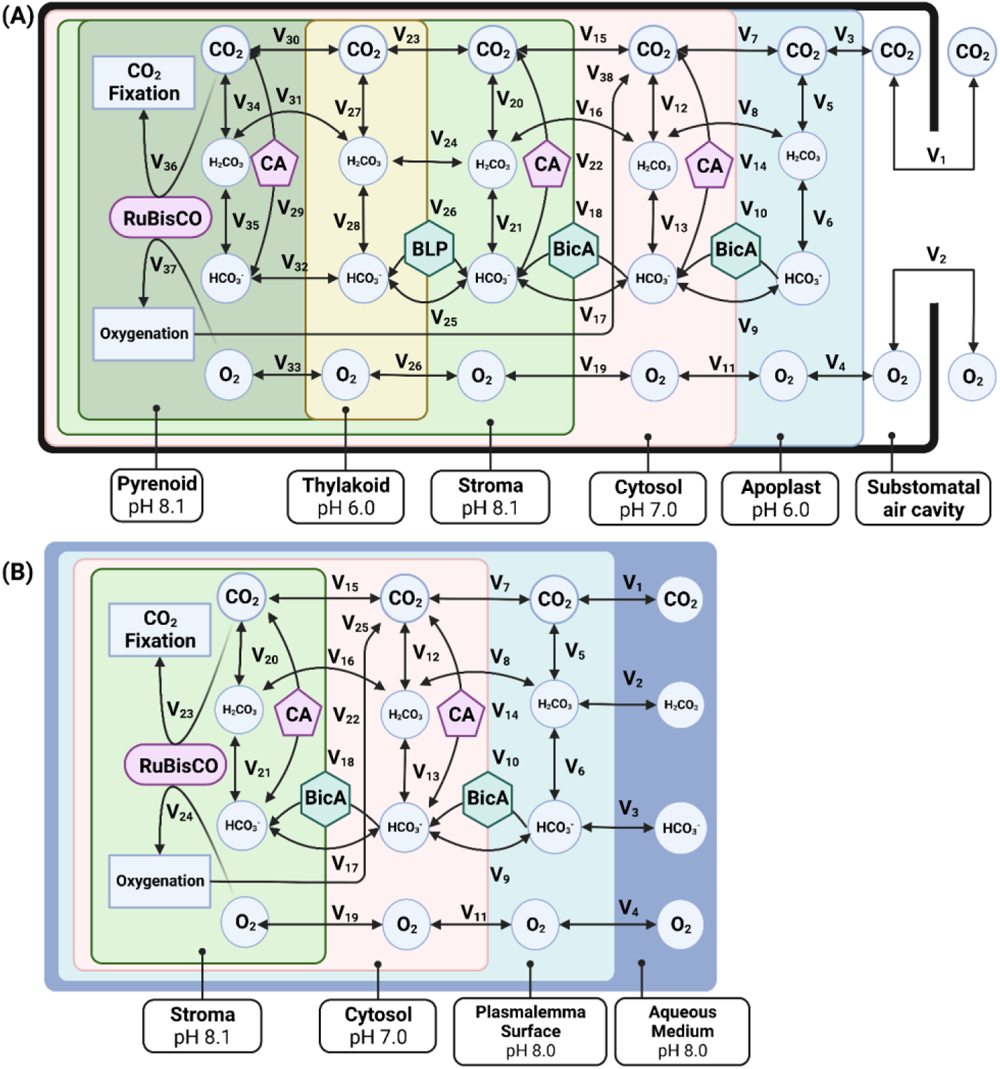
Diagrammatic representations of (A) a model of photosynthetic carbon assimilation in a land plant mesophyll cell containing a *C. reinhardtii* style PCCM, and (B) a model of an algal cell that does not contain a pyrenoid. CA refers to carbonic anhydrase, BLP refers to bestrophin-like proteins that serve as membrane channels for passive bicarbonate transport, and BicA is a cyanobacterial active bicarbonate transporter. In the *VCell* implementation of the model, some strongly linked steps are combined for the sake of numerical computability. Exact specifications for all flux equations used can be found in the publicly shared model implementations in *VCell* (see code and data availability statement). Note that for the sake of numerical tractability, the carbonic-anhydrase catalyzed interconversion of CO_2_ and HCO_3_ in the thylakoid in models featuring a CCM (v29) is localized to the pyrenoid but uses the pH value of the thylakoid; in the real biological system, the carbonic-anhydrase is inside the thylakoid tubules that penetrate into the pyrenoid.

**Table 1:** Model parameter definitions with source references and, where applicable with notes on derivation. When parameters were derived from parameterization of a previous modeling study, both the modeling study and the original literature reference for the parameter are cited. References in “Ref.” column: (1) Mazarei & Sandall 1980; (2) Fei *et al*. 2022; (3) Xiang & Anderson 1994; (4) Walker, Smith & Cathers 1980; (5) Bentley & Pittman 1997; (6) Gutknecht, Bisson & Tosteson 1977; (7) Missner *et al*. 2008; (8) Hopkinson *et al*. 2011; (9) Widomska, Raguz & Subczynski 2007; (10) Mitchell *et al*. 2010; (11) Larsson *et al*. 1997; (12) Pocker & Ng 1973; (13) Pocker & Miksch 1978; (14) McGrath & Long 2014; (15) Bernacchi *et al*. 2005; (16) Badger & Andrews 1974; (17) Farquhar, von Caemmerer & Berry 1980; (18) von Caemmerer 2000; (19) Price *et al*. 2004; (20) Bernacchi *et al*. 2006; (21) Kump 2008; (22) Pritchard, Grout & Short 1986; (23) Flexas *et al*. 2021; (24) Ouk, Oi & Taniguchi 2020; (25) Slaton & Smith 2002; (26) Yu, Tang & Kuo (2000); (27) Feely, Doney & Cooley (2009); (28) Felle 2001; (29) Kramer, Sacksteder & Cruz 1999.

| Name | Value(s) | Units | Notes | Ref. |
| --- | --- | --- | --- | --- |
| Diffusion coefficient of CO<sub>2</sub> in water | 1.88 x 10<sup>3</sup> | um<sup>2</sup> s<sup>-1</sup> |  | (1, 2) |
| Diffusion coefficient of H<sub>2</sub>CO<sub>3</sub> in water | 1.2 x 10<sup>3</sup> | um<sup>2</sup> s<sup>-1</sup> | Assumed in (Fei *et al.*, 2022) to be identical to diffusion coefficient of acetic acid | (2, 3) |
| Diffusion coefficient of HCO<sub>3</sub><sup>-</sup> in water | 1.15 x 10<sup>3</sup> | um<sup>2</sup> s<sup>-1</sup> |  | (2, 4) |
| Diffusion coefficient of O<sub>2</sub> in water | 2.42 x 10<sup>3</sup> | um<sup>2</sup> s<sup>-1</sup> |  | (5) |
| Membrane permeability to CO<sub>2</sub> | 3.50e-03;<br>3.20e-02 | m s<sup>-1</sup> | Parameter scanned between reported values | (6, 7) |
| Membrane permeability to H<sub>2</sub>CO<sub>3</sub> | 30 | um s<sup>-1</sup> |  | (2) |
| Membrane permeability to HCO<sub>3</sub><sup>-</sup> | 0.05 | 0.05 um s<sup>-1</sup> |  | (8) |
| Membrane permeability to O<sub>2</sub> | 75 | cm s<sup>-1</sup> |  | (9) |
| Besotrophin-like channel mediated permeability of thylakoid to HCO<sub>3</sub><sup>-</sup> | 1 x 10<sup>-2</sup> | m s<sup>-1</sup> |  | (2) |
| Chloroplast membrane permeability to HCO<sub>3</sub><sup>-</sup> mediated by LCIA | 1 x 10<sup>-8</sup> | m s<sup>-1</sup> |  | (2) |
| Rate constant of spontaneous hydration of CO<sub>2</sub> | 6 x 10<sup>-2</sup> | s<sup>-1</sup> |  | (10) |
| Rate constant of spontaneous dehydration of H<sub>2</sub>CO<sub>3</sub> | 2 x 10<sup>1</sup> | s<sup>-1</sup> |  | (10) |
| Rate constant of spontaneous deprotonation of H<sub>2</sub>CO<sub>3</sub> | 1 x 10<sup>7</sup> | s<sup>-1</sup> |  | (10) |
| Rate constant of spontaneous protonation of HCO<sub>3</sub><sup>-</sup> | 5 x 10<sup>10</sup> | M<sup>-1</sup> s<sup>-1</sup> |  | (10) |
| Carbonic anhydrase k<sub>cat</sub> | 0.3 x 10<sup>6</sup> | s<sup>-1</sup> |  | (11) |
| Carbonic anhydrase K<sub>m</sub> for CO<sub>2</sub> | 1.5 | mol m<sup>-3</sup> |  | (12) |
| Carbonic anhydrase K<sub>eq</sub> | 0.56 x 10<sup>-6</sup> | mol m<sup>-3</sup> |  | (13) |
| Carbonic anhydrase K<sub>m</sub> for HCO<sub>3</sub><sup>-</sup> | 34 | mol m<sup>-3</sup> |  | (13) |
| Carbonic anhydrase concentration in stroma | 270 | uM |  | (14) |
| Carbonic anhydrase concentration in cytosol | 135 | uM | Assumed to be half the stroma value | (14) |
| Carbonic anhydrase concentration in lumen | 135 | uM | Assumed to be half the stroma value | (14) |
| Rubisco V<sub>max</sub> of carboxylation | 7600 | umol L<sup>-1</sup> s<sup>-1</sup> |  | (15) |
| Rubisco V<sub>max</sub> of oxygenation | 1596 | umol L<sup>-1</sup> s<sup>-1</sup> | Calculated from ratio of k<sub>cat</sub> values of carboxylation and oxygenation. | (16, 17) |
| Rubisco K<sub>m</sub> O<sub>2</sub> | 8.6 | umol L<sup>-1</sup> |  | (18) |
| Rubisco K<sub>m</sub> CO<sub>2</sub> | 215 | umol L<sup>-1</sup> |  | (18) |
| BicA V<sub>max</sub> | 1.85 x 10<sup>-4</sup> | mol m<sup>-2</sup> s<sup>-1</sup> | Parameter scanned | (19) |
| BicA K<sub>m</sub> HCO<sub>3</sub><sup>-</sup> | 0.217 | mol m<sup>-3</sup> |  | (19) |
| Stomatal conductance | 0.4375 | mol m<sup>-2</sup> s<sup>-1</sup> |  | (20) |
| Atmospheric concentration of CO<sub>2</sub> | 412 | ppm |  | Assumed |
| Atmospheric concentration of O<sub>2</sub> | 0.21 | Partial pressure |  | (21) |
| Thickness of cell wall in angiosperms | 0.32 | um |  | (14, 22) |
| Thickness of cell wall in bryophytes | 1.6 | um |  | (23) |
| Effective porosity of C3 plant cell wall | 0.2 | Unitless |  | (14) |
| Effective porosity of hornwort cell wall | 0.0001 |  | Parameter scanned | Calculated (23) |
| Thickness of unstirred boundary layer in algal model | 0.32 | um | Assumed to be the same as cell wall thickness | Assumed |
| Thickness of unstirred apoplast water layer in land plant models | 0.32 | um | Assumed to be the same as cell wall thickness | Assumed |
| Permeability of pyrenoid starch sheath to dissolved inorganic carbon | 0.1 * P<sub>CO2</sub> | um s<sup>-1</sup> | From range of permeabilities that allow effective carbon concentration in modeling done by (Hopkinson et al., 2011) | (2) |
| Permeability of pyrenoid starch sheath to oxygen | 0.1 * P<sub>O2</sub> | um s<sup>-1</sup> | Assumed to behave similarly to dissolved inorganic carbon | Assumed |
| Radius of pyrenoid | 1.0 | um |  | (2) |
| Radius of thylakoid | 0.5 | um | Multiplied by 10X to account for simpler thylakoid architecture | (2) |
| Height of thylakoid | 4 | um |  | Assumed |
| Radius of chloroplast | 4.63 | um | Calculated from stroma volume fraction and assuming spherical geometry | (14) |
| Radius of cytosol | 8.77 | um | Assuming spherical geometry | (24) |
| Radius of plasmalemma surface | 9.23 | um | Calculated from radius of cytosol, cell wall thickness, and assumed apoplast water thickness | Calculated |
| Radius of substomatal space in land plant model | 11.63 | um |  | (14) |
| Proportion of cell wall adjacent to intercellular airspace in land plant | 0.5 | Unitless |  | (25) |
| pH of land plant apoplast | 6.0 | pH |  | (26) |
| pH of ocean water | 8.1 | pH |  | (27) |
| pH of cytosol | 7.2 | pH |  | Calculated (28) |
| pH of stroma | 8.0 | pH |  | (29) |
| pH of lumen | 6.0 | pH |  | (29) |

Systems were represented as spatially symmetrical, with spherical concentric compartments that were converted into volumetric pixels (voxels) according to the simulations’ spatial resolution. All results presented are from simulations containing either 9,261 voxels or 12,167 voxels. Due to the large parameter explorations done in this study, minor geometrical modifications were made to make efficient numerical simulation feasible. Specifically, the radius of the apoplast water layer in the land plant models was extended out from the 9.41um it should be based on a cell wall thickness of 0.32um plus an apoplast water layer of equivalent thickness to 10um. We also modeled the thylakoid tubules of with-PCCM models as a set of six cylinders of radius 0.5um extending into the pyrenoid, with exchange between the tubules and the pyrenoid occurring at the end of these cylinders, in contrast to the larger number of finer tubules used in (Fei et al., 2022).

### Reaction equations

Carboxylation flux by rubisco is calculated as in (Farquhar et al., 1980) **(E1)**. The rate of carboxylation by rubisco is normally taken to be the minimum of V_c_ and J, where J describes the rate of ribulose-1,5-bisphosphate regeneration enabled by photosynthetic electron transport and a function of J_max_, a maximum rate of RuBP regeneration, among other parameters (Farquhar et al., 1980). Estimates of the relevant parameters are available for land plants but, to our knowledge, not for algae. We are also specifically examining CO_2_-limiting conditions where rubisco reaction rate limitations dominate. For these reasons, we calculate the carboxylation and oxygenation rates assuming that the system is not limited by RuBP regeneration as in (Fei et al., 2022).

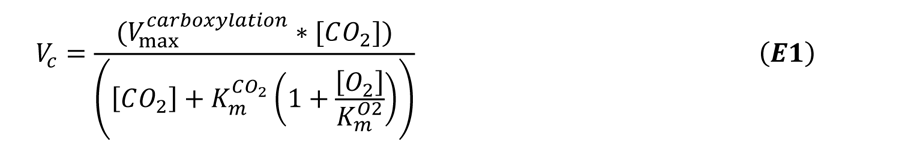

The ratio of oxygenation to carboxylation Vmax is:

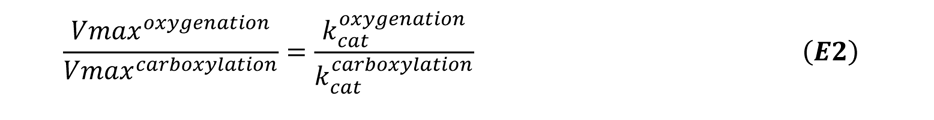

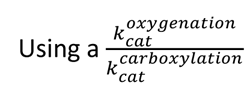 value of 0.21 as in (Farquhar et al., 1980), we can thereby calculate the *Vmax^oxygenation^* of our systems. The oxygenation flux by rubisco is then calculated as:

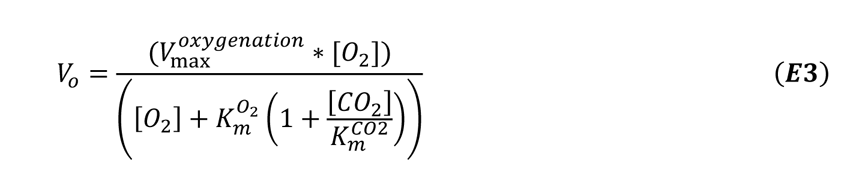

Interconversion of CO_2_ with bicarbonate via carbonic anhydrase is described as in (McGrath and Long, 2014):

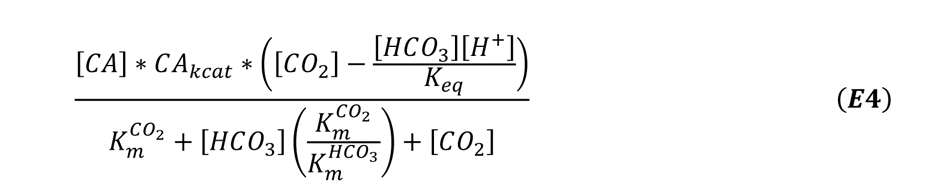

In the land plant models, the flux density of dissolution of gaseous CO_2_ or O_2_ into the water layer is as in (Hemond and Fechner, 2022):

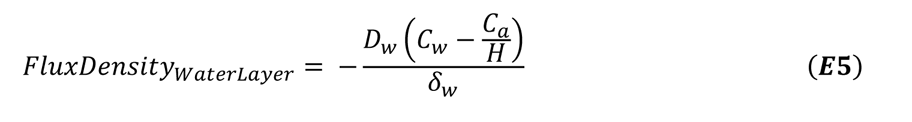

Where *D_w_* is the diffusion rate of the dissolving species in water *C_w_ and C_a_* are the concentrations of that species in the air and in the water layer, *H* is the dimensionless Henry’s Law constant, and ^δ^ is the length of the unstirred water layer into which the gas is dissolving. In our models, we assume the presence of a thin layer of water on top of the plant’s cell wall that is the same thickness as the cell wall itself into which CO_2_ is dissolving.

Permeation of aqueous species through the cell wall is given by the following equation, as in (McGrath and Long, 2014):

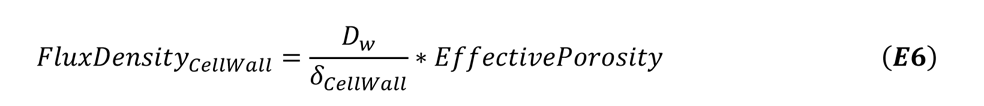

Where *EffectivePorosity* is the porosity of the cell wall divided by the tortuosity of the cell wall.

For computational tractability, we combine the processes of gases dissolving into water and the aqueous species passing through the cell wall. Note that in the above equation *D_w_* / δ_*w*_ and *D_w_ *EffectivePorosity*/δ_CellWall_ gives permeability (in units of um/s) of the water layer and the cell wall, respectively. Multiplying these values by surface area gives conductivities (in units of µm^3^/s). The inverses of these values are resistances, which can be summed to give the total resistance of the water layer plus the cell wall. The inverse of this, again, will be the conductivity of the overall system, which can be multiplied by the concentration gradient from the air to the surface of the plasmalemma to give the total flux.

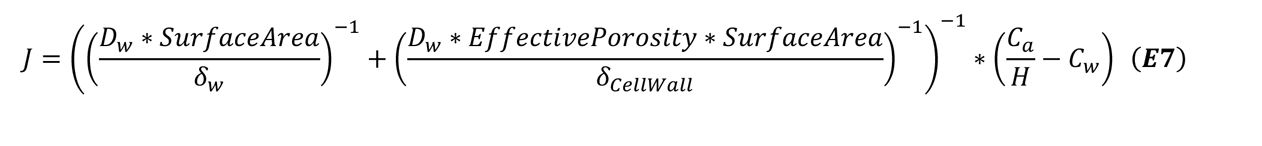

Permeation through lipid membranes is given by:

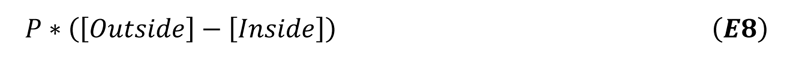

Active transport by bicarbonate transporter *BicA* is described using Michaelis-Menten kinetics:

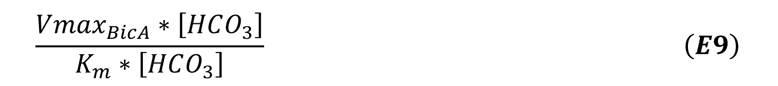

*Efficiency Calculations*

Net CO_2_ fixation is described as:

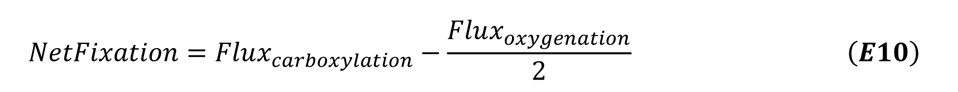

2 NADPH equivalents are expended per carboxylation or oxygenation reaction based on the stoichiometry of the CBC cycle and photorespiration. 3 ATP and 3.5 ATP are used for a single carboxylation or oxygenation event, respectively (Edwards and Walker, 1983).

In models featuring a PCCM, there is a lumenal carbonic anhydrase that catalyzes the following reaction:

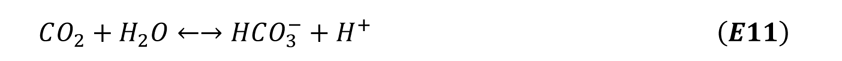

Due to the acidic pH of the lumen (Kramer et al., 1999) the net flux of this reaction is overwhelmingly in the direction of CO_2_ and H_2_O, so that entry of bicarbonate depletes the proton motive force (pmf) that is maintained by the light reactions of photosynthesis, which imposes an indirect ATP cost on CCM activity by requiring additional proton pumping to maintain the pmf (Mukherjee et al., 2019). Based on a 14:3 ratio of pumped protons to ATP synthesis via the thylakoid membrane ATP synthase, inferred from the number of c-subunits in such ATP synthases (Seelert et al., 2000), we can calculate the indirect ATP cost of this lumen CA activity as:

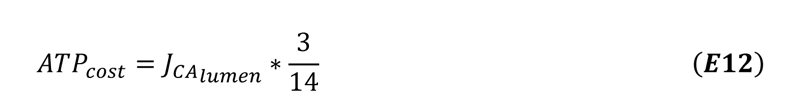

This is added to the other ATP consumption in the model (due to the metabolic costs of carboxylation and oxygenation) to give total ATP use. This can be compared with NADPH use due to carboxylation and oxygenation to get an estimate of the total ATP, NADPH, and the ATP:NADPH ratio needed to support the activity in the model. From the values provided in (Walker et al., 2020) we estimate the amount of either Cyclic Electron Flow (CEF) or Malate Valve activity needed to rebalance the ATP/NADPH ratio needed for a particular model, which we can then convert into an additional demand for photons and, therefore, a the number of photons needed on a per reaction (carboxylation or oxygenation) basis (**Figure S1**). From this, we can calculate the number of photons needed to support model fluxes and then compare this to the net fixation achieved by a model to get an estimate of light use efficiency.

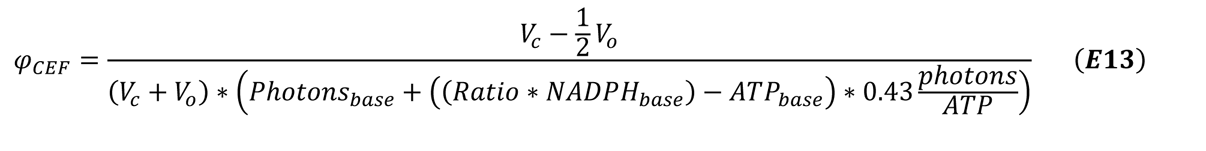

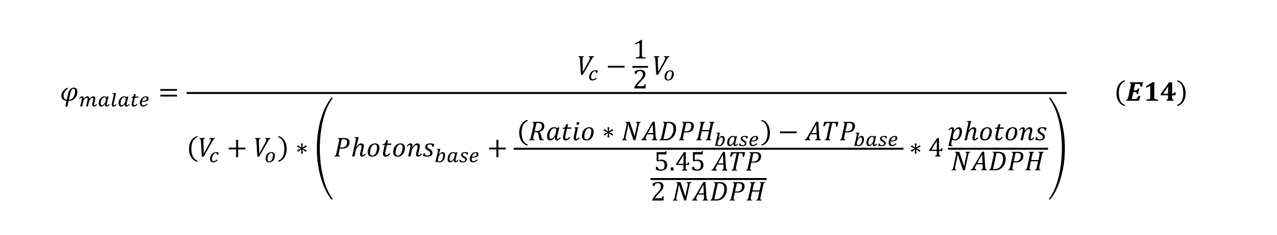

Where *Vc and Vo* are the modeled rates of carboxylation and oxygenation, *Ratio* refers to the modeled ATP/NADPH ratio necessary to support the fluxes in the model, and Photons_base_, ATP_base_ and NADPH_base_ refer to the photons used and the ATP and NADPH generated in the process of making two NADPH molecules via Linear Electron Flow (LEF) (Walker et al., 2020).

### Concentration calculations

All concentrations in the models used in this study are in units of µM. To calculate the µM concentrations of CO_2_ and O_2_ in the atmosphere, we used the following conversion:

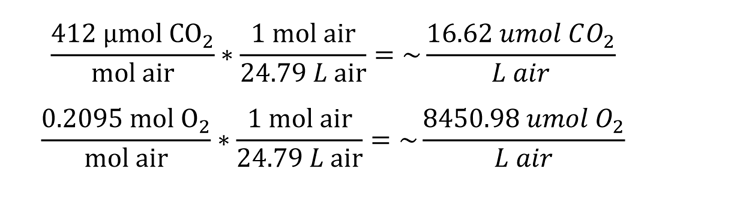

## Results

### Validation of compensation point predictions and sensitivity analysis

The land plant and algal carbon assimilation models were validated by comparing a key estimated result (CO_2_ compensation point) with experimentally measured values from the literature. The CO_2_ compensation point is the external CO_2_ level at which net CO_2_ assimilation by a photosynthesizing organism is zero (i.e. carbon assimilation by rubisco is balanced out by CO_2_ losses to photorespiration and respiration in the light, denoted as R_L_). Low compensation points are also a defining feature of organisms with CCMs since they maintain net positive carbon assimilation at lower CO2 concentrations, making this a useful indicator of whether land plant and algal models with and without CCMs reasonably recreate the carbon assimilation dynamics of real systems.

As shown in Figure 2 and **Table 2**, the models with CCMs have substantially lower compensation points than the models lacking CCMs. Moreover, as shown in **Table 2**, these estimated compensation point values fall within the ranges of values reported in the literature for angiosperm land plants and algae with and without CCMs (Table 2). Note that the reported compensation points of hornworts with pyrenoids (11-13 ppm) are lower than those of closely related C3 liverworts, but higher than typical estimates for C4 plants and pyrenoid-containing algae (Villarreal and Renner, 2012).

**Figure 2:**
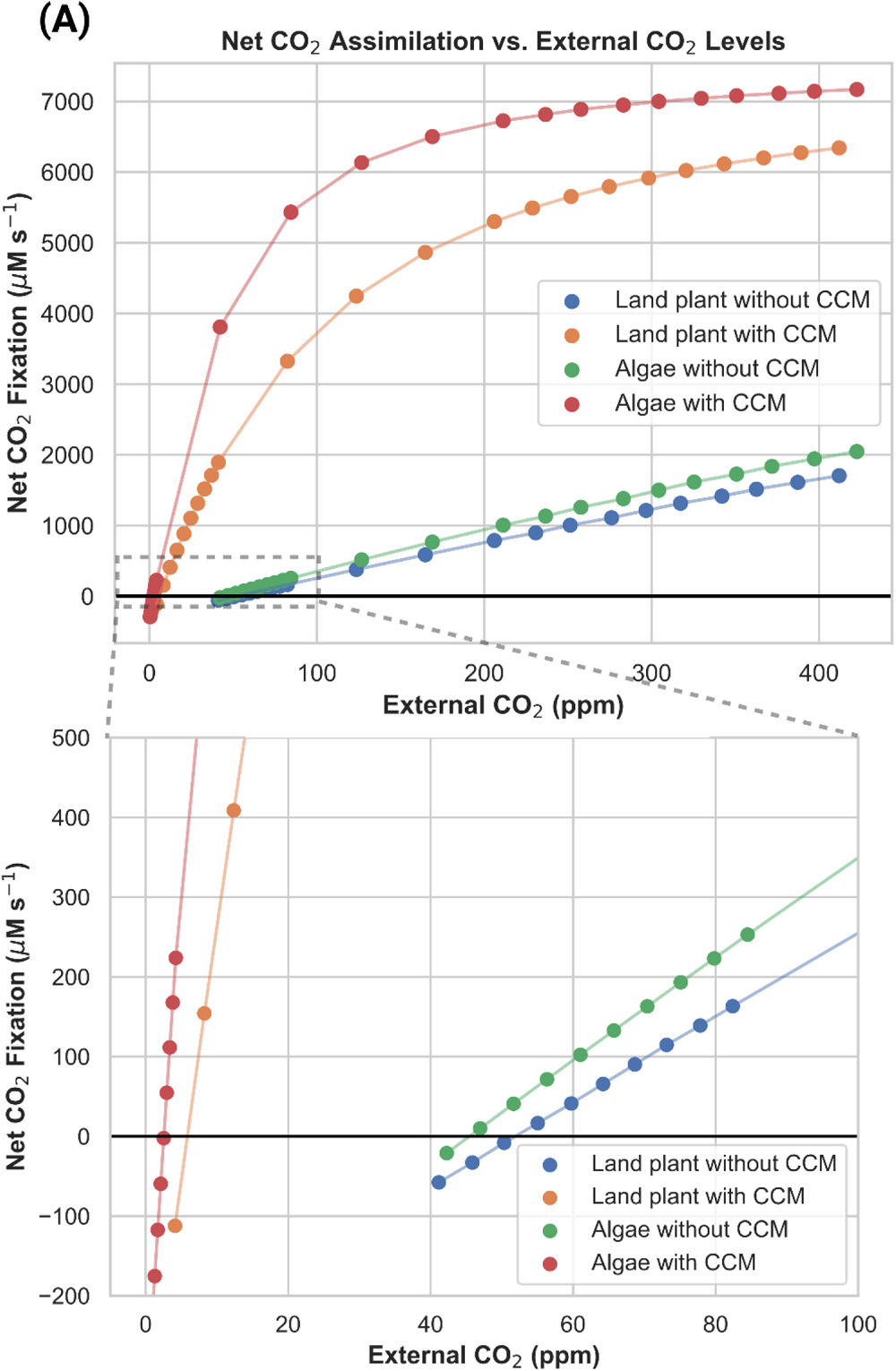
Net CO_2_ assimilation versus external CO_2_ concentrations in carbon assimilation models. The point at which net CO_2_ assimilation is zero defines the compensation point. **(A)** The full range of saturation and external CO_2_ concentrations, and **(B)** a zoomed-in panel showing the point at which each curve reaches 0% rubisco saturation (i.e. the compensation point).

**Table 2:** Predicted compensation points for different models from the present study compared with reference values from the literature. Reference column numbers refer to their numbering in the bibliography.

| Model | Compensation point<br>(ppm CO<sub>2</sub>) | Reference Values<br>(ppm CO<sub>2</sub>) | Measurement<br>References |
| --- | --- | --- | --- |
| Land plant with CCM | 6.2 | 1.3; 4.3; 0.7 – 9.0 | (Fladung and Hesselbach, 1987; Lee et al., 2022) |
| Land plant without CCM | 52.7 | 48; 57; 48.2 – 53.4; 65-100 | (Fladung and Hesselbach, 1987; Tolbert et al., 1995; Peixoto et al., 2021) |
| Algal model with CCM | 2.7 | 0.75 – 2.5; 6.0 | (Coleman and Colman, 1980; Raven et al., 1982) |
| Algal model without CCM | 44.6 | 43.5 – 58; 64.5 | (Raven et al., 1982; Steensma et al., 2023) |

The sensitivity analysis results shown in Figure 3 show that simulated net CO_2_ assimilation and quantum yield values from the land plant models are relatively robust to local variations in all parameters, providing us with confidence that these results are not merely the result of a very particular selection of parameters. In both the land plant and algal models without PCCMs, rubisco V_max_, cell and chloroplast radii, and membrane permeability to CO_2_ are the most influential determinants of net CO_2_ assimilation and quantum yield. In the land plant model, stomatal conductance also stands out. The addition of a PCCM reduces the sensitivity of net CO_2_ assimilation to changes in any input parameter but increases the sensitivity of the predicted quantum yield to input parameter values. The local stability of our results to perturbations in key parameters is comparable with previous studies, being more variable than the models presented in (Fei et al., 2022), which spatially modeled a smaller system (algal chloroplasts), and significantly less variable than the models presented in (McGrath and Long, 2014), which modeled land plant CO_2_ assimilation at a similar scale. We also characterized the sensitivity of our modeling results to the spatial resolution of the numerical simulations. Our results **(Figure S2-3)** show that rubisco saturation - the percentage of maximum rubisco activity achieved – and quantum yield in an algal model lacking a CCM are robust to the simulation resolution. Increasing the resolution all the way down to 0.32um, well beyond what could feasibly be done given the amount of parameter exploration done in this study, does result in noticeable changes in pyrenoid [CO_2_] and [HCO_3+_], resulting in small increases in rubisco saturation and small decreases in quantum yield **(Figure S4-5)**.

**Figure 3:**
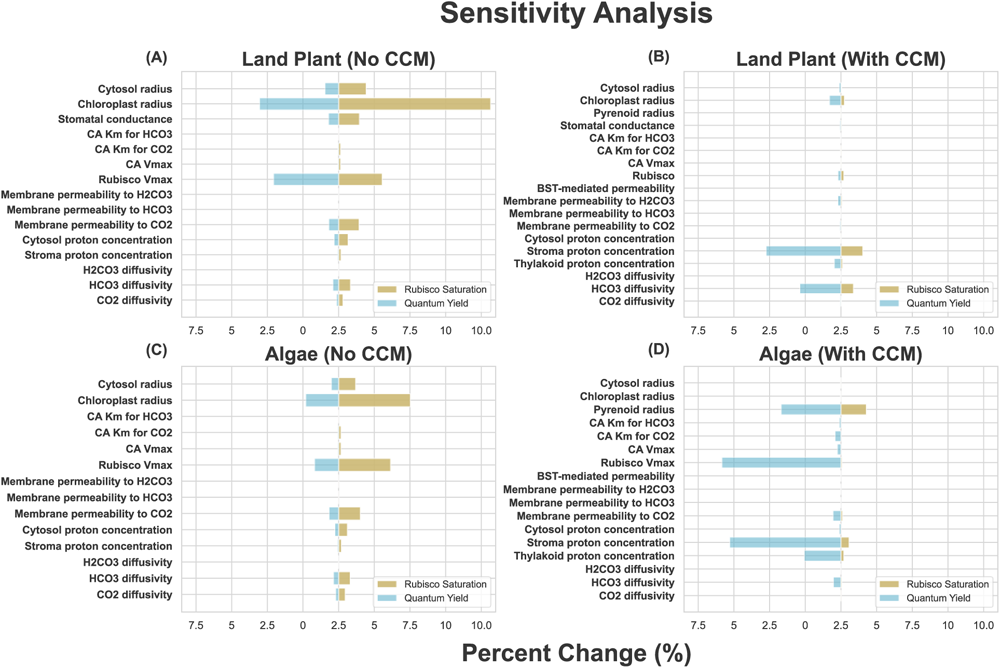
Sensitivity analysis results for **(A)** the land plant model lacking a CCM, **(B)** the algal model lacking a CCM, **(C)** the land plant model with a CCM, and **(D)** the algal model with a CCM. Orange bars indicate the absolute % change of quantum yield resulting from a 10% change in the indicated parameter, and blue bars represent the same for rubisco saturation. For both of the land plant models, increasing the cytosol radius by 10% resulted in problems with solving the systems numerically, so the cytosol radius was increased by 1% instead and, assuming a linear relationship between the size of radius increase and the change in rubisco saturation and quantum yield, multiplied by 10 to get the values shown in **(A-B)**.

### Efficiency of chloroplast membrane bicarbonate channel is strongly dependent on assumed permeability of chloroplast membrane to CO_2_

Previous studies (Price et al., 2010; McGrath and Long, 2014) have suggested that the incorporation of bicarbonate transporters into the chloroplast membrane of a land plant could improve net fixation and/or the efficiency of carbon assimilation, and that this could represent a reasonable intermediate stage in a broader biotechnological effort to implement a full CCM in a land plant. Modeling studies on CCM systems typically assume the lipid membrane permeability of 0.35 cm/s, which was experimentally measured and reported in (Gutknecht et al., 1977). However, there is substantial uncertainty as to the value of parameter, with experimental estimates ranging over many orders of magnitude (Evans et al., 2009). The permeability may be as much as an order of magnitude higher than the Gutknecht *et al* value, as reported in (Missner et al., 2008). We hypothesized that the apparent favorability of employing a chloroplast membrane bicarbonate pump may be highly sensitive to the assumed chloroplast membrane CO_2_ permeability.

To test this hypothesis, we performed a parameter exploration from an order of magnitude lower than the widely cited (Gutknecht et al., 1977) value up to the (Missner et al., 2008) value in both land plant and algal systems, calculating net fixation as well as ATP/CO_2_ and light-use efficiency, as shown in Figure 4.

**Figure 4:**
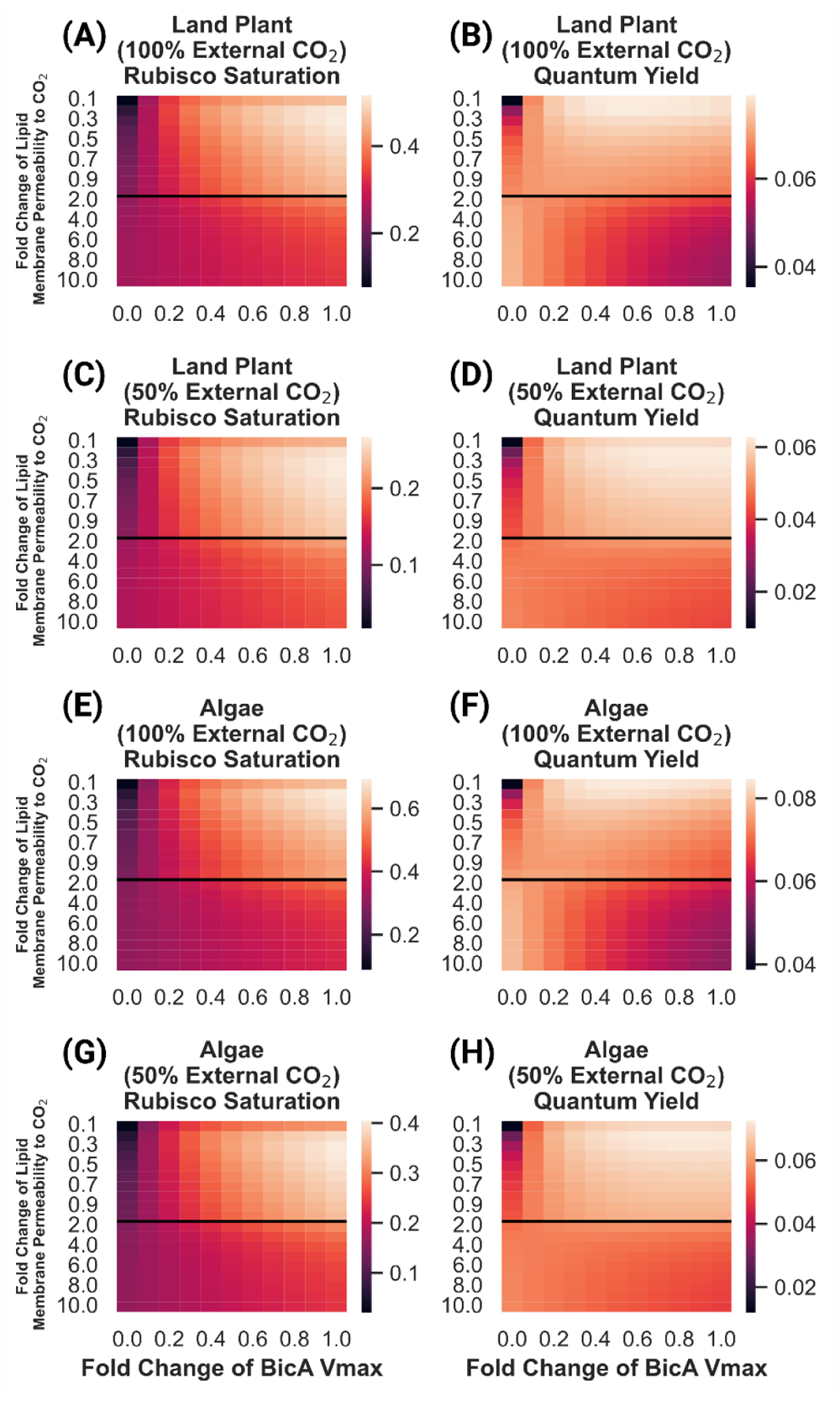
Rubisco saturation and quantum yield of land plant and algal models of CO_2_ assimilation under 100% and 50% external CO_2_ levels, as a function of lipid membrane permeability to CO_2_ and BicA bicarbonate transporter V_max_. Fold change of lipid membrane permeability is relative to the value reported in (Gutknecht et al., 1977). **(A)** Predicted rubisco saturation of a land plant model under 100% external CO_2_. **(B)** Predicted quantum yield of a land plant model under 100% external CO_2_. **(C)** Predicted rubisco saturation of a land plant model under 50% external CO_2._ **(D)** Predicted quantum yield of a land plant model under 50% external CO_2_. **(E)** Predicted rubisco saturation of an algal model under 100% external CO_2._ **(F)** Predicted quantum yield of an algal model under 100% external CO_2_. **(G)** Predicted rubisco saturation of an algal model under 50% external CO_2._ **(H)** Predicted quantum yield of an algal model under 50% external CO_2_. The black lines in each plot indicate the (Gutknecht et al., 1977) value for lipid bilayer permeability to CO_2_ as well as a transition in the y-axis from increments of 0.1X to 1X fold changes.

These results show that the light use efficiency of a chloroplast membrane bicarbonate transporter is highly sensitive to the value of the chloroplast envelope’s permeability to CO_2_, with a large range of permeabilities resulting in 2X more ATP usage per unit of CO_2_ fixed. In the land plant model, we see increases in both rubisco saturation and quantum yield as BicA pumping activity increases when lipid membrane permeability values are equivalent to, or below that reported in (Gutknecht et al., 1977) (Figure 4A-B). At permeabilities higher than this, increased BicA activity actually decreases quantum yield, though net fixation still increases (Figure 4A-B). We see a similar picture in the algal model (Figure 4E-F), suggesting that the differences in DIC form, concentration, and diffusivity do not greatly impact the sensitivity of this strategy to the specific value of lipid membrane permeability to CO_2._ The decrease in quantum yield in models with high lipid membrane permeability to CO_2_ is driven by increased leakage of CO_2_ from the chloroplast back into the cytosol after it interconverts with the bicarbonate just pumped by BicA **(shown as flux V_15_ in** Figure 1). As lipid membranes become more permeable to CO_2_, its tendency to escape the chloroplast before being fixed by rubisco increases. Lowering the external CO_2_ concentration does, however, change the energy efficiency penalty of increased BicA activity significantly (**Figure 4C-D;G-H**). Even at higher lipid membrane permeability values, we see only minimal decreases in quantum yield with increased BicA bicarbonate pumping.

### Efficiency of a plasmalemma bicarbonate channel is strongly dependent on external DIC levels and limited by the rate of equilibration between CO_2_ and bicarbonate

We found that although the strategy of pumping bicarbonate from the cytosol to the chloroplast may incur substantial energy costs, implementing a bicarbonate pump at the plasmalemma may be more effective. This makes sense considering that in aqueous systems at near-neutral pH, most of the DIC in the system is in the form of bicarbonate. We incorporated a plasmalemma bicarbonate transporter and explored the efficiency of such a system across different external DIC concentrations and activities of the transporter in both algal and land plant systems (Figure 5).

**Figure 5:**
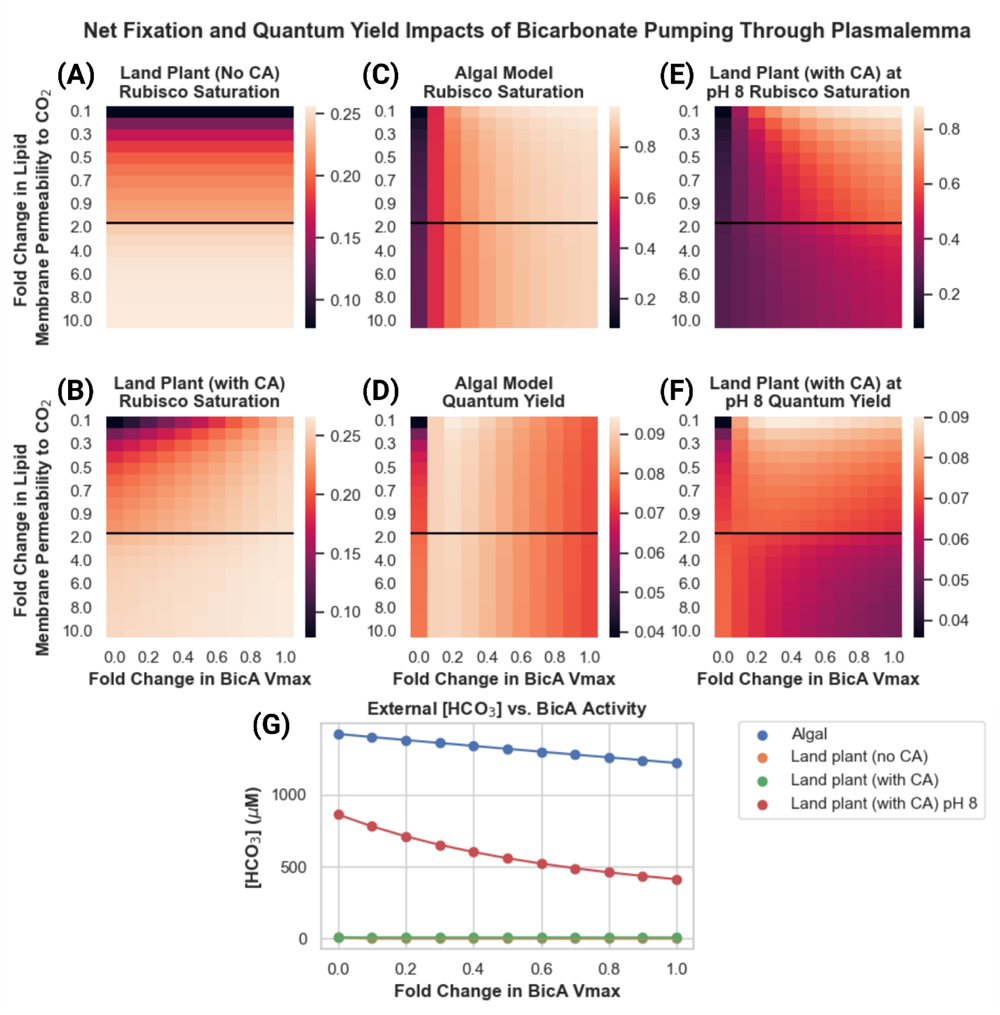
Predicted rubisco saturation and quantum yield in land plant and algal models with a BicA bicarbonate pump present in the plasmalemma membrane, as a function of assumed lipid membrane permeability to CO_2_ and BicA V_max_. Fold change of lipid membrane permeability is relative to the value reported in (Gutknecht et al., 1977). **(A)** Predicted rubisco saturation of a land plant model lacking an apoplastic carbonic anhydrase. **(B)** Predicted rubisco saturation of a land plant model with an apoplastic carbonic anhydrase. **(C)** Predicted rubisco saturation of an algal model. **(D)** Predicted quantum yield of an algal model. **(E)** Predicted rubisco saturation of a land plant model with an apoplastic carbonic anhydrase and an apoplast pH of 8. **(F)** Predicted quantum yield of a land plant model with an apoplastic carbonic anhydrase and an apoplast pH of 8. The black lines in each plot indicate the (Gutknecht et al., 1977) value for lipid bilayer permeability to CO_2_ as well as a transition in the y-axis from increments of 0.1X to 1X fold changes.

In the land plant model, the plasmalemma bicarbonate pump is not an effective means of increasing either net fixation or energy efficiency. As anticipated, the pump does work in the algal case (Figure 5). The key difference appears to be that the external environment in the algal system, which is suffused with bicarbonate ions, can maintain reasonably high steady-state concentrations in the vicinity of the cell to support the bicarbonate pumping activity (Figure 5C-D). In contrast, in the land plant system all dissolved bicarbonate available to the cell must first enter the system as CO_2_ in the intercellular airspace, dissolve into the water in the apoplast, and then spontaneously hydrate to H_2_CO_3_ and deprotonate into bicarbonate. Although the protonation/deprotonation between H_2_CO_3_ is extremely fast, the hydration/dehydration is not (first-order rate constant of hydration of CO_2_ to H_2_CO_3_ is 6 x 10^-2^ s^-1^ (Mitchell et al., 2010)). The result is an almost instantaneous depletion of the HCO_3-_ concentration in the apoplast space, with insufficient spontaneous hydration flux to replenish it (Figure 5G). Adding carbonic anhydrase activity to the apoplast allows for much faster regeneration of the external HCO_3-_ concentration, allowing BicA to impact rubisco saturation (Figure 5A-B). However, the pH of the apoplast, although variable, tends to be slightly to moderately acidic (Yu et al., 2000), resulting in low HCO_3-_ concentrations in the land plant model even with the apoplast carbonic anhydrase included (Figure 5G). It is only when the apoplast pH is made substantially more basic (pH of 8) and a carbonic anhydrase is included that the land plant model can replicate the algal model’s rubisco saturation and quantum yield gains by using a plasmalemma bicarbonate pump (Figure 5E-F).

### PCCM integration results in greater marginal cost of CO_2_ fixation improvements in land plants vs. algal systems and switches from decreasing to increasing light-use efficiency around a C_i_ typical of C4 plants

We compared the energy-use efficiency of PCCM integration by comparing the predicted cost in photons of fixing CO_2_ molecule in four different models: (i) a land plant model with a PCCM, (ii) a land plant model without a PCCM, (iii) an algal model with a PCCM, and (iv) an algal model without a PCCM. By dividing the increase in net CO_2_ fixation in models (i) and (iii) relative to models (ii) and (iv) we estimated the marginal cost of in photons of fixing an additional CO_2_ molecule using a PCCM in our land plant and models (Figure 6). As we observed when examining the efficiency of the plasmalemma and chloroplast envelope BicA bicarbonate pumps, the assumed permeability of lipid membranes can have an impact on efficiency; in this case, however, the relative marginal cost values do not change dramatically between an assumed permeability equivalent to that used in previous studies (1.0 in Figure 6A-B) and the higher value closer to that reported in (Missner et al., 2008).

**Figure 6:**
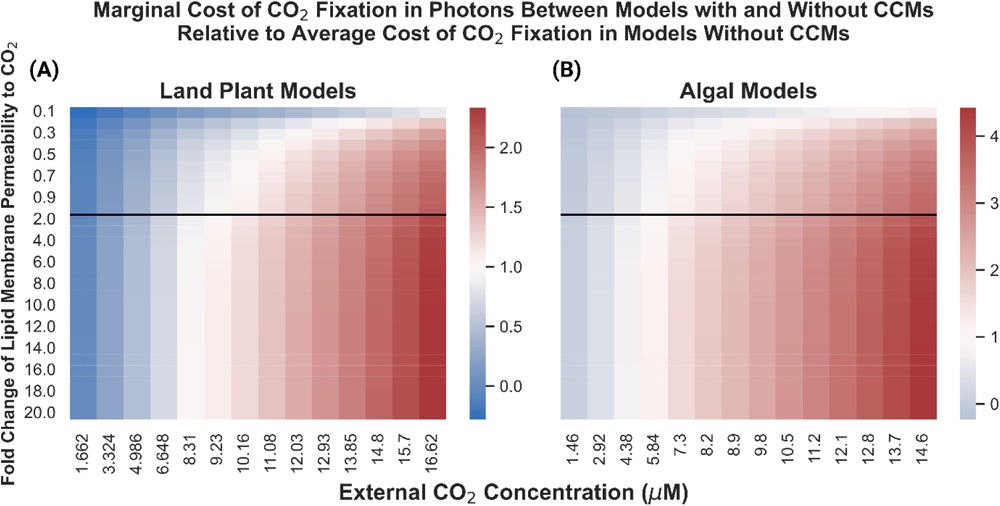
The ratio of the marginal cost in photons of one unit of net CO_2_ fixation in land plant **(A)** and algal **(B)** models resulting from adding a PCCM relative to the average cost of fixing one molecule of CO_2_ in those same models without CCMs, as a function of lipid membrane permeability and external CO_2_ concentrations. Fold change of lipid membrane permeability is relative to the value reported in (Gutknecht et al., 1977). Blue indicates that for a given lipid membrane permeability / external CO_2_ concentration combination, the model containing a CCM has a lower marginal cost of CO_2_ fixation – ie., is more light-efficient – than the average cost of CO_2_ fixation in the model lacking a CCM. Red indicates that for a given parameterization, the model containing a CCM has a higher marginal cost of CO_2_ fixation than the average cost of CO_2_ fixation in its CCM lacking counterpart. The black lines in each plot indicate the (Gutknecht et al., 1977) value for lipid bilayer permeability to CO_2_ as well as a transition in the y-axis from increments of 0.1 to 1 in the X-fold changes.

In the algal models, the use of the PCCM appears to only become marginally efficient with respect to light usage below an external [CO_2_] of 4.38 uM. In contrast, the CCM is efficient in the land plant model below a substomatal [CO_2_] of 243 ppm.

#### As cell wall thickness increases and cell wall effective porosity decreases, PCCMs become more favorable in land plant models

Given the findings regarding PCCMs in land plants highlighted above, it is interesting that many species of hornworts have pyrenoids – are there any meaningful biophysical differences between hornworts and other land plants that could explain these differences? As highlighted in (Meyer et al., 2008; Flexas et al., 2021) hornworts and other bryophytes have cell walls that are both substantially thicker and less porous compared to other land plants. From the mesophyll conductance values reported for angiosperms and bryophytes reported in (Meyer et al., 2008; Flexas et al., 2021), and with the assumption that other internal resistances to CO_2_ diffusion are similar between bryophytes and embryophytes, we can estimate that the effective porosity of a bryophyte like a hornwort must be on the order of four orders of magnitude smaller than in a typical C3 angiosperm. We explore parameters within this range of possible porosity values and across multiple external CO_2_ concentrations (Figure 7).

**Figure 7:**
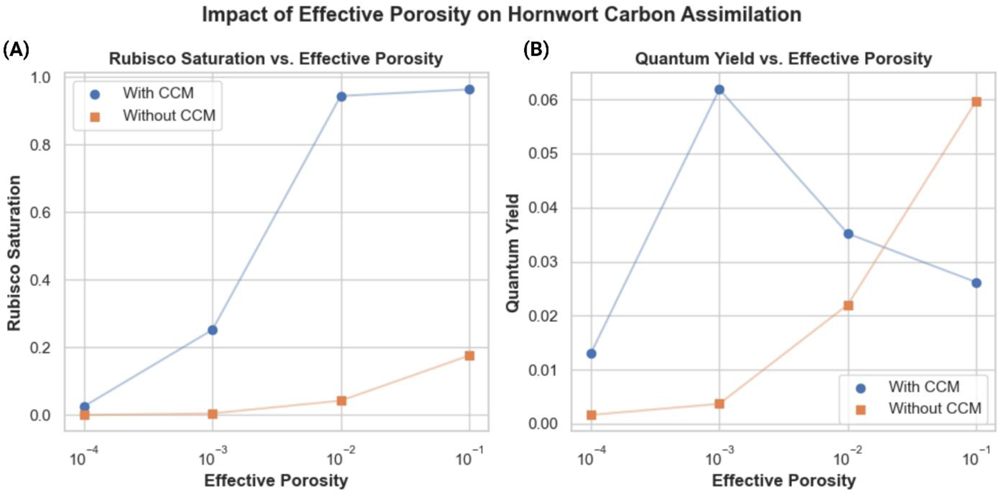
Rubisco saturation **(A)** and quantum yield **(B)** of a land plant model with varying effective porosity values. Blue points / lines represent predicted rubisco saturation or quantum yield in models including a PCCM; orange points/ lines represent predicted saturation or quantum yield in models not including a PCCM.

Below effective porosities on the order of 10^-1^, which fall in the range we would expect of angiosperms, our model shows that the plant struggles to fix CO_2_ without a CCM. With a PCCM, however, the model can achieve some level of net CO_2_ fixation all the way down to effective porosities of 10^-3^. Below porosities of 10^-3^, we do not observe net CO_2_ fixation in the model without a PCCM, and at a porosity of 10^-4^, both models with and without PCCMs struggle to fix carbon. In terms of light-use efficiency, the model with a PCCM achieves a greater quantum yield of photosynthesis than the model without a PCCM below effective porosities of 10^-2^.

## Discussion

We initially hypothesized that the conspicuous absence of biophysical CCMs in almost all land plant lineages, in contrast to algae where they are widespread (Raven et al., 2005), may be the result of lower efficiency of such systems in land plants relative to algae, and that this results from their different biophysical contexts. To our surprise, we found that PCCMs appear to result in qualitatively similar improvements in quantum yield and net CO_2_ assimilation in land plant and algal models. In the algal model, the fact that addition of a PCCM does not result in efficiency gains until relatively low external DIC levels are reached is surprising, given that *Chlamydomonas reinhardtii* cells appear to concentrate carbon even at recent “air-level” – approximately 330ppm – CO_2_ concentrations (Badger et al., 1980). This implies that algae may routinely run their CCMs even when this incurs a quantum yield penalty. In contrast, the intercellular CO_2_ concentration at which the CCM improves quantum yield in the land plant model (∼243ppm) is higher than reported estimates of C_i_ in C4 plants under laboratory, greenhouse, and field conditions (Bunce, 2005). Previous work has described the evolutionary history of C4 photosynthesis (Sage et al., 2018) and identified certain anatomical features – namely Kranz anatomy – and environmental factors such as hot, arid conditions that lead to increased transpirational water loss and factors such as Water-Use Efficiency (WUE) as key predictors of C4 emergence. If the estimated quantum yield gains resulting from the introduction of a biophysical CCM to a land plant in this study apply to biochemical CCMs like C4 and CAM photosynthesis, this may represent an additional evolutionary driver towards such systems.

Hornworts are the only land plant lineage that has evolved a biophysical CCM and they have done so multiple times (Villarreal and Renner, 2012). Hornworts, as well as some other bryophytes, are noteworthy for having substantially slower gas exchange between their surroundings and their photosynthetic tissues when compared with vascular land plants (Meyer et al., 2008).. Our results show that a land plant with the low effective cell wall porosities we might expect given their extremely poor gas exchange characteristics, the use of a CCM becomes necessary to achieve net CO_2_ fixation, which would impose a strong selective pressure for adopting one. The fact that hornworts represent the earliest-diverging extant branch of the land plants, and therefore may have maintained the genes and regulatory networks necessary to adopt a PCCM, may explain why this biophysical CCM strategy has been adopted by hornworts and not other land plants growing in conditions where biochemical CCMS have been selected for. We should note that in the models presented in this study, at effective porosities below 10^-3^, only single digit values of rubisco saturation are achieved even with a biophysical CCM present and active, which may not be sufficient for viability, especially since we do not have or include estimates of respiration in the light in the models. This is despite the fact that mesophyll conductance to CO_2_ in hornworts, which we are using effective porosity as a proxy for in this study, has been measured to be four-to-five orders of magnitude lower than in angiosperms (Flexas et al., 2021). This suggests that our model underestimates the strength of the hornwort CCM or otherwise does not properly describe some aspect of hornwort CO_2_ assimilation. The ratio of chloroplast-to-thallus surface area has not been explored in our modeling, but was found in a previous study to be a potentially important determinant of hornwort mesophyll conductance (Carriquí et al., 2019). Future work might aim to incorporate an exploration of chloroplast position and surface area to better account for this in the modeling.

These results shed light on potential challenges associated with improving crop productivity via the introduction of biophysical CCMs. The specific value chosen for the permeability of lipid bilayers to CO_2_ has a large effect on the predicted energy efficiency of our models, with values higher than those used in previous modeling studies (McGrath and Long, 2014; Fei et al., 2022) but within the range of previously reported literature values (Gutknecht et al., 1977; Missner et al., 2008) resulting in qualitatively different conclusions. We see this in our consideration of BicA-mediated HCO3^-^ pumping, which had been previously flagged as a promising intermediate step in introducing a biophysical CCM to a C3 plant (Price et al., 2010; McGrath and Long, 2014). As noted in (Fei et al., 2022), barriers to CO_2_ diffusion form a key component of known functional CCMs, so the finding that the chloroplast membrane may provide enough of a diffusion barrier for the transport of HCO3^-^ into the stroma and subsequent conversion to CO_2_ to meaningfully improve net fixation and carbon assimilatory efficiency was surprising. Our results show that at or below the permeability reported in (Gutknecht et al., 1977), which is used in other modeling studies, increasing BicA pumping activity leads to improvements in quantum yield, indicating more efficient CO_2_ fixation with respect to light use. However, above this value, we see uniform decreases in quantum yield with increased BicA activity. Net CO_2_ fixation increases with BicA pumping in all cases; therefore, in situations where light is abundant relative to CO_2_, this decrease in efficiency may not impact plant fitness. However, recent modeling work suggests that *J_max_*, the maximum rate of ribulose-1,5-bisphosphate (RuBP) regeneration enabled by photosynthetic electron transport, is more limiting to crop yield than limits to the maximum rate of carboxylation (V_max_ of rubisco carboxylation) under the projected elevated atmospheric CO_2_ levels of 2050 and 2100 (He and Matthews, 2023). In this study, improved quantum yields correspond to a combination of (i) lower expenditures of ATP for each CO_2_ molecule fixed, and (ii) a more favorable ATP/NADPH ratio needed for fixation, resulting in less energy loss from the use of Cyclic Electron Flow during ATP/NADPH rebalancing (Walker et al., 2020). Under conditions of *J_max_* limitations, differences in quantum yield may become a critical factor in determining yield, making the sensitivity of quantum yield in this and other studies to assumed lipid bilayer permeability to CO_2_ a matter of critical importance.

Interestingly, previous studies in this area (McGrath and Long, 2014; Fei et al., 2022) have performed sensitivity analyses that include this permeability as a surveyed parameter and its modeled effect is small compared to other parameters. These small local sensitivity values are estimated by observing the change in an output value like light-saturated CO_2_ assimilation with a ± 10% change in the permeability parameter. This ignores the fact that the uncertainty in this value is in the range of at least an order of magnitude (Evans et al., 2009), and so despite low local sensitivity, the overall change that can result from varying it within reasonable bounds is substantial. The substantial uncertainty in this critical parameter could be reined in by future experimental measurements, though this will still be complicated by the potentially large variation between different plant systems, dynamic remodeling of lipid bilayers in response to developmental and environmental cues, etc. In the absence of well-defined values for this parameter, we encourage future groups modeling such systems to explore a range of values and to characterize the robustness of their conclusions to its variation.

In the near-neutral or slightly basic conditions that most photosynthetic organisms in aqueous environments find themselves in, HCO_3-_ represents the primary form of Dissolved Inorganic Carbon (DIC) in their surroundings. Due to the impermeability of lipid bilayers to passive diffusion of HCO_3_ ^-^, the use of this pool of DIC requires organisms to employ an active transport mechanism (e.g. cyanobacterial HCO_3_ ^-^ pumps like BicA (Price et al., 2004)) to move it from the extracellular to the intracellular space, which may often make sense due to the sheer quantity of DIC that is present in the environment. Although land plants ultimately obtain CO_2_ from the atmosphere, this CO_2_ must dissolve into water prior to entering photosynthesizing cells, at which point this aqueous CO_2_ interconverts with other DIC species. This raises the possibility of a similar strategy – pumping HCO_3_ ^-^ from a land plant’s apoplast water into the intracellular environment to increase net CO_2_ fixation – potentially viable. However, our results indicate that the limited spontaneous rate of CO_2_ and HCO_3_ ^-^ interconversion without the activity of carbonic anhydrase means that this strategy does not work.

Of note here is the fact that a quantitatively very similar system arises in algae growing in acidic environments where external HCO_3_ ^-^ levels are negligible, such as the red alga *Cyanidioschyzon merolae* (De Luca et al., 1978). In such systems, all DIC must first enter the cell passively as aqueous CO_2_, at which point it will interconvert primarily between CO_2_ and HCO_3_, with the ratio of CO_2_:HCO_3_ determined by the cytosolic pH. There is strong evidence that *C. merolae* has a non-pyrenoid based CCM (Steensma et al., 2023). Such a system could use HCO_3_ ^-^ pumping across the chloroplast envelope as a method of concentrating carbon, but our results suggest that this system would require maintenance of a near-neutral cytosolic pH along with the presence of carbonic anhydrases in the cytosol to be viable. The maintenance of this near-neutral pH in an acidic environment may, in turn, represent a substantial energetic cost to the organism.

## Data and Code Availability

All results generated as part of this study can be found in the Supplemental Material. Models used for generating the results can all be found under the account *kastejos* in the Virtual Cell interface. Specific model names can be found for each dataset in the corresponding Supplemental Material tables.

## Acknowledgments

The Virtual Cell, the software platform used for the reaction-diffusion simulations in this study, is supported by NIH Grant R24 GM137787.

This research was supported by the U.S. Department of Energy, Office of Science Biological and Environmental Research Grant no DE-SC0018269 (J.A.M.K., Y.S-H.) and Basic energy Sciences Grant no DE-FG02-91ER20021 (B.J.W.). This work is supported, in part, by the NSF Research Traineeship Program (Grant DGE-1828149) to J.A.M.K. This publication was also made possible by a predoctoral training award to J.A.M.K. from Grant T32-GM110523 from National Institute of General Medical Sciences (NIGMS) of the NIH. Its contents are solely the responsibility of the authors and do not necessarily represent the official views of the NIGMS or NIH.

## Author Contributions

J.A.M.K, B.J.W, and Y.S-H. conceptualized the study. J.A.M.K. developed the models, ran the simulations, and analyzed the results. J.A.M.K. wrote the first draft of the manuscript. All authors contributed to revising and editing the final manuscript.

## References

1. Badger MR, Kaplan A, Berry JA (1980) Internal inorganic carbon pool of Chlamydomonas reinhardtii: evidence for a carbon dioxide-concentrating mechanism. Plant physiology 66: 407–413

2. Bräutigam A, Schlüter U, Eisenhut M, Gowik U (2017) On the Evolutionary Origin of CAM Photosynthesis. Plant Physiol 174: 473–477

3. Bunce J (2005) What is the usual internal carbon dioxide concentration in C 4 species under midday field conditions? Photosynthetica 43: 603–608

4. Carriquí M, Roig-Oliver M, Brodribb TJ, Coopman R, Gill W, Mark K, Niinemets Ü, Perera-Castro AV, Ribas-Carbó M, Sack L, et al (2019) Anatomical constraints to nonstomatal diffusion conductance and photosynthesis in lycophytes and bryophytes. New Phytologist 222: 1256–1270

5. Coleman JR, Colman B (1980) Effect of oxygen and temperature on the efficiency of photosynthetic carbon assimilation in two microscopic algae. Plant Physiol 65: 980–983

6. Cowan AE, Moraru II, Schaff JC, Slepchenko BM, Loew LM (2012) Spatial modeling of cell signaling networks. Methods in cell biology 110: 195–221

7. De Luca P, Taddei R, Varano L (1978) Cyanidioschyzon merolae: a new alga of thermal acidic environments. Webbia 33: 37–44

8. Edwards G, Walker D (1983) C3, C4: Mechanisms, Cellular and Environmental Regulation of Photosynthesis. Univ of California Press

9. Ermakova M, Danila FR, Furbank RT, von Caemmerer S (2020) On the road to C(4) rice: advances and perspectives. Plant J 101: 940–950

10. Evans JR, Kaldenhoff R, Genty B, Terashima I (2009) Resistances along the CO2 diffusion pathway inside leaves. Journal of Experimental Botany 60: 2235–2248

11. Farquhar GD, Caemmerer S, Berry JA (1980) A biochemical model of photosynthetic CO2 assimilation in leaves of C3 species. Planta 149: 78–90

12. Fei C, Wilson AT, Mangan NM, Wingreen NS, Jonikas MC (2022) Modelling the pyrenoid-based CO2-concentrating mechanism provides insights into its operating principles and a roadmap for its engineering into crops. Nature Plants 8: 583–595

13. Fladung M, Hesselbach J (1987) Developmental Studies on Photosynthetic Parameters in C3, C3–C4 and C4 Plants of Panicum. Journal of Plant Physiology 130: 461–470

14. Flexas J, Clemente-Moreno MJ, Bota J, Brodribb TJ, Gago J, Mizokami Y, Nadal M, Perera-Castro AV, Roig-Oliver M, Sugiura D, et al (2021) Cell wall thickness and composition are involved in photosynthetic limitation. Journal of experimental botany 72: 3971–3986

15. Gutknecht J, Bisson MA, Tosteson FC (1977) Diffusion of carbon dioxide through lipid bilayer membranes: effects of carbonic anhydrase, bicarbonate, and unstirred layers. The Journal of general physiology 69: 779–794

16. He Y, Matthews ML (2023) Seasonal climate conditions impact the effectiveness of improving photosynthesis to increase soybean yield. Field Crops Research 296: 108907

17. Hemond HF, Fechner EJ (2022) Chemical fate and transport in the environment. Academic Press

18. Hennacy JH, Jonikas MC (2020) Prospects for Engineering Biophysical CO(2) Concentrating Mechanisms into Land Plants to Enhance Yields. Annu Rev Plant Biol 71: 461–485

19. Hopkinson BM, Dupont CL, Allen AE, Morel FMM (2011) Efficiency of the CO2-concentrating mechanism of diatoms. Proceedings of the National Academy of Sciences 108: 3830–3837

20. Kramer DM, Sacksteder CA, Cruz JA (1999) How acidic is the lumen? Photosynthesis Research 60: 151– 163

21. Lee M-S, Boyd RA, Ort DR (2022) The photosynthetic response of C3 and C4 bioenergy grass species to fluctuating light. GCB Bioenergy 14: 37–53

22. Ludwig M (2013) Evolution of the C4 photosynthetic pathway: events at the cellular and molecular levels. Photosynth Res 117: 147–161

23. McGrath JM, Long SP (2014) Can the cyanobacterial carbon-concentrating mechanism increase photosynthesis in crop species? A theoretical analysis. Plant physiology 164: 2247–2261

24. Meyer M, Seibt U, Griffiths H (2008) To concentrate or ventilate? Carbon acquisition, isotope discrimination and physiological ecology of early land plant life forms. Philosophical transactions of the Royal Society of London Series B, Biological sciences 363: 2767–2778

25. Missner A, Kügler P, Saparov SM, Sommer K, Mathai JC, Zeidel ML, Pohl P (2008) Carbon dioxide transport through membranes. The Journal of biological chemistry 283: 25340–25347

26. Mitchell MJ, Jensen OE, Cliffe KA, Maroto-Valer MM (2010) A model of carbon dioxide dissolution and mineral carbonation kinetics. Proceedings of the Royal Society A: Mathematical, Physical and Engineering Sciences 466: 1265–1290

27. Mukherjee A, Lau CS, Walker CE, Rai AK, Prejean CI, Yates G, Emrich-Mills T, Lemoine SG, Vinyard DJ, Mackinder LCM, et al (2019) Thylakoid localized bestrophin-like proteins are essential for the CO(2) concentrating mechanism of Chlamydomonas reinhardtii. Proc Natl Acad Sci U S A 116: 16915–16920

28. Peixoto MM, Sage TL, Busch FA, Pacheco HDN, Moraes MG, Portes TA, Almeida RA, Graciano-Ribeiro D, Sage RF (2021) Elevated efficiency of C3 photosynthesis in bamboo grasses: A possible consequence of enhanced refixation of photorespired CO2. GCB Bioenergy 13: 941–954

30. Price GD, Badger MR, von Caemmerer S (2010) The Prospect of Using Cyanobacterial Bicarbonate Transporters to Improve Leaf Photosynthesis in C3 Crop Plants. Plant Physiology 155: 20–26

31. Price GD, Woodger FJ, Badger MR, Howitt SM, Tucker L (2004) Identification of a SulP-type bicarbonate transporter in marine cyanobacteria. Proceedings of the National Academy of Sciences 101: 18228–18233

32. Raven JA, Ball LA, Beardall J, Giordano M, Maberly SC (2005) Algae lacking carbon-concentrating mechanisms. Can J Bot 83: 879–890

33. Raven JA, Beardall J, Johnston AM (1982) Inorganic Carbon Transport in Relation to H+ Transport at the Plasmalemma of Photosynthetic Cells. Plasmalemma and Tonoplast: Their Functions in the Plant Cell. Elsevier Biomedical Press, Amsterdam, pp 41–47

34. Raven JA, Beardall J, Sánchez-Baracaldo P (2017) The possible evolution and future of CO2-concentrating mechanisms. Journal of Experimental Botany 68: 3701–3716

35. Raven JA, Cockell CS, De La Rocha CL (2008) The evolution of inorganic carbon concentrating mechanisms in photosynthesis. Philos Trans R Soc Lond B Biol Sci 363: 2641–2650

36. Sage RF, Monson RK, Ehleringer JR, Adachi S, Pearcy RW (2018) Some like it hot: the physiological ecology of C(4) plant evolution. Oecologia 187: 941–966

37. Schaff J, Fink CC, Slepchenko B, Carson JH, Loew LM (1997) A general computational framework for modeling cellular structure and function. Biophysical journal 73: 1135–1146

38. Seelert H, Poetsch A, Dencher NA, Engel A, Stahlberg H, Müller DJ (2000) Proton-powered turbine of a plant motor. Nature 405: 418–419

39. Steensma AK, Shachar-Hill Y, Walker BJ (2023) The carbon-concentrating mechanism of the extremophilic red microalga Cyanidioschyzon merolae. Photosynth Res 156: 247–264

40. Still CJ, Berry JA, Collatz GJ, DeFries RS (2003) Global distribution of C3 and C4 vegetation: Carbon cycle implications. Global Biogeochemical Cycles 17: 6–1

41. Tolbert NE, Benker C, Beck E (1995) The oxygen and carbon dioxide compensation points of C3 plants: possible role in regulating atmospheric oxygen. Proceedings of the National Academy of Sciences 92: 11230–11233

42. Villarreal JC, Renner SS (2012) Hornwort pyrenoids, carbon-concentrating structures, evolved and were lost at least five times during the last 100 million years. Proceedings of the National Academy of Sciences 109: 18873–18878

43. Walker BJ, Kramer DM, Fisher N, Fu X (2020) Flexibility in the Energy Balancing Network of Photosynthesis Enables Safe Operation under Changing Environmental Conditions. Plants (Basel, Switzerland). doi: 10.3390/plants9030301

44. Walker BJ, VanLoocke A, Bernacchi CJ, Ort DR (2016) The Costs of Photorespiration to Food Production Now and in the Future. Annu Rev Plant Biol 67: 107–129

45. Yu Q, Tang C, Kuo J (2000) A critical review on methods to measure apoplastic pH in plants. Plant and Soil 219: 29–40

