## Supplemental Material for "Biophysical carbon concentrating mechanisms in land plants: insights from reaction-diffusion modeling"

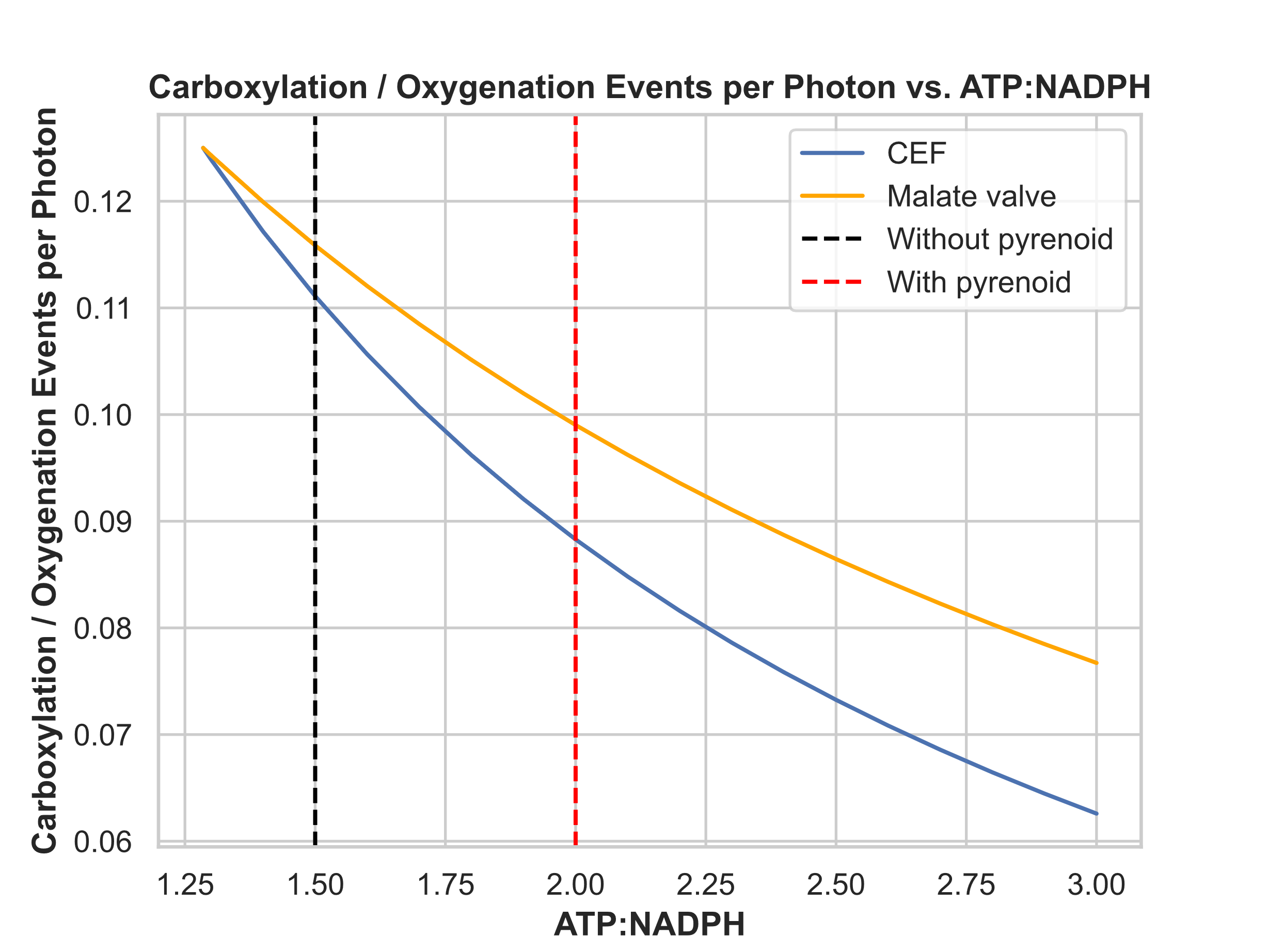

**Figure S1:** Carboxylation / oxygenation events per photon as a function of varying ATP:NADPH ratios. Costs associated with using either Cyclic Electron Flow or the malate valve for increasing ATP:NADPH ratio from the products of the light reactions are taken from (1).

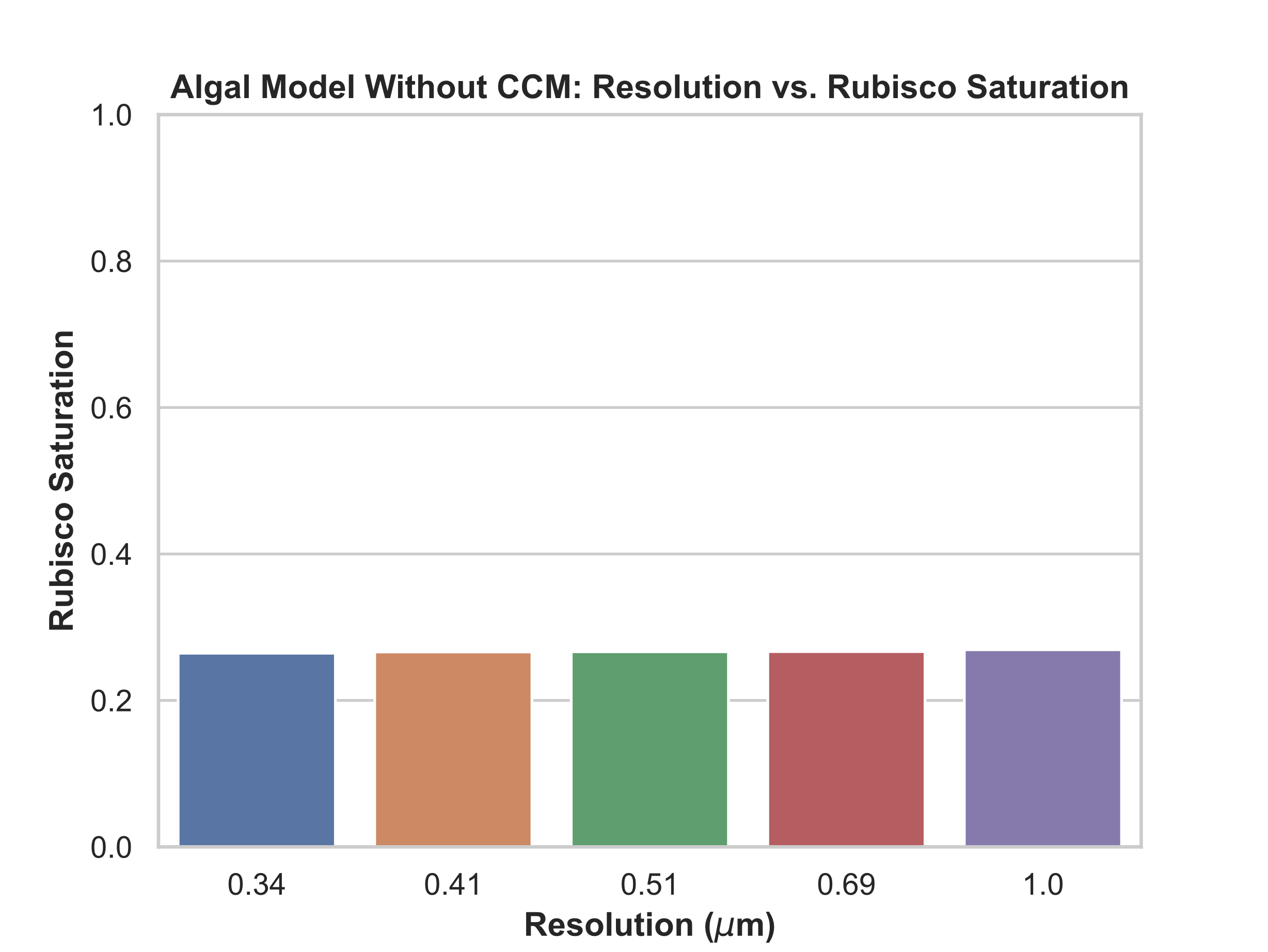

**Figure S2:** Effect of simulation spatial resolution on rubisco saturation. Simulation results are taken from the model of an algal cell without a CCM.

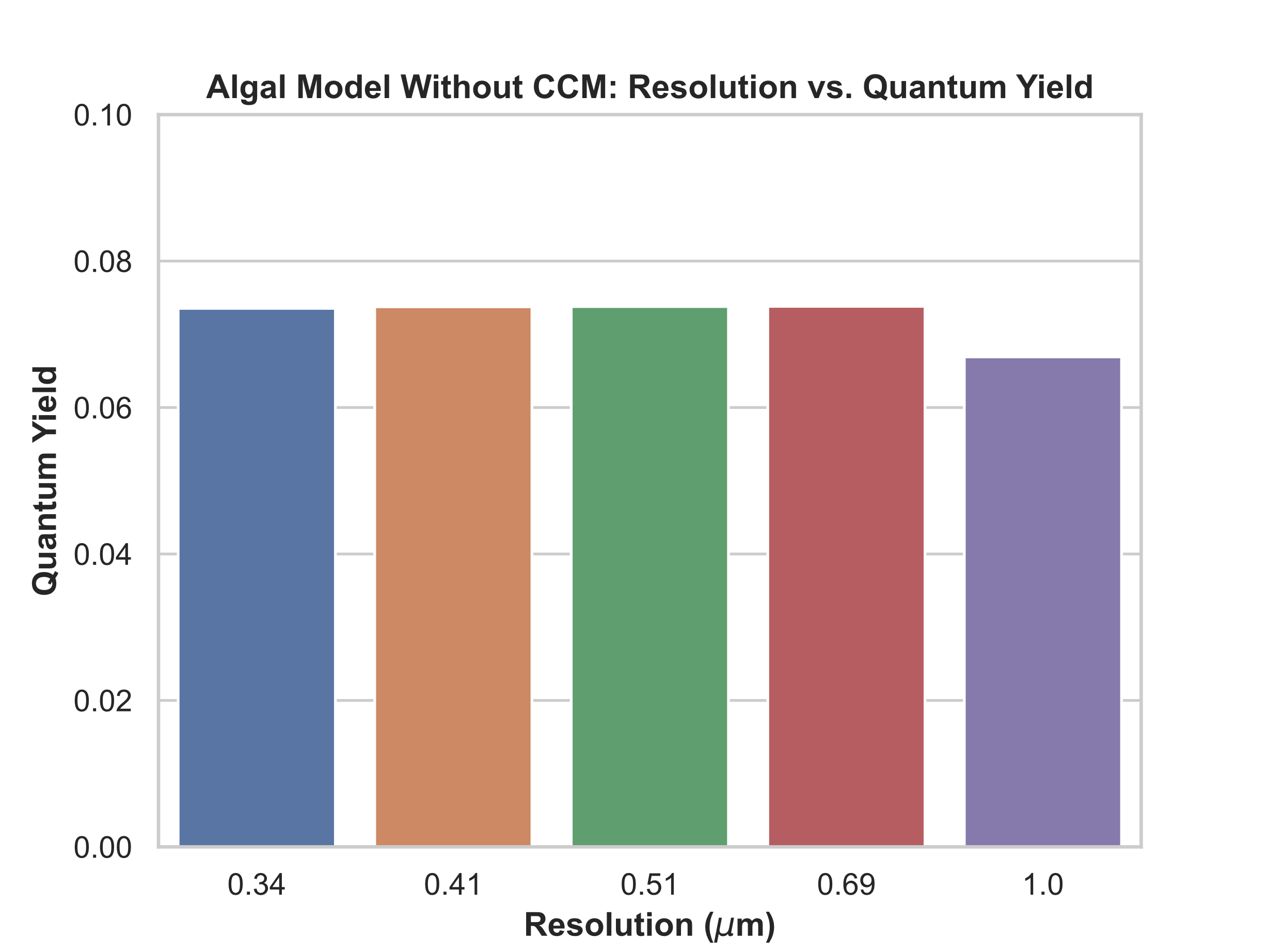
 **Figure S3:** Effect of simulation spatial resolution on quantum yield. Simulation results are taken from the model of an algal cell without a CCM.

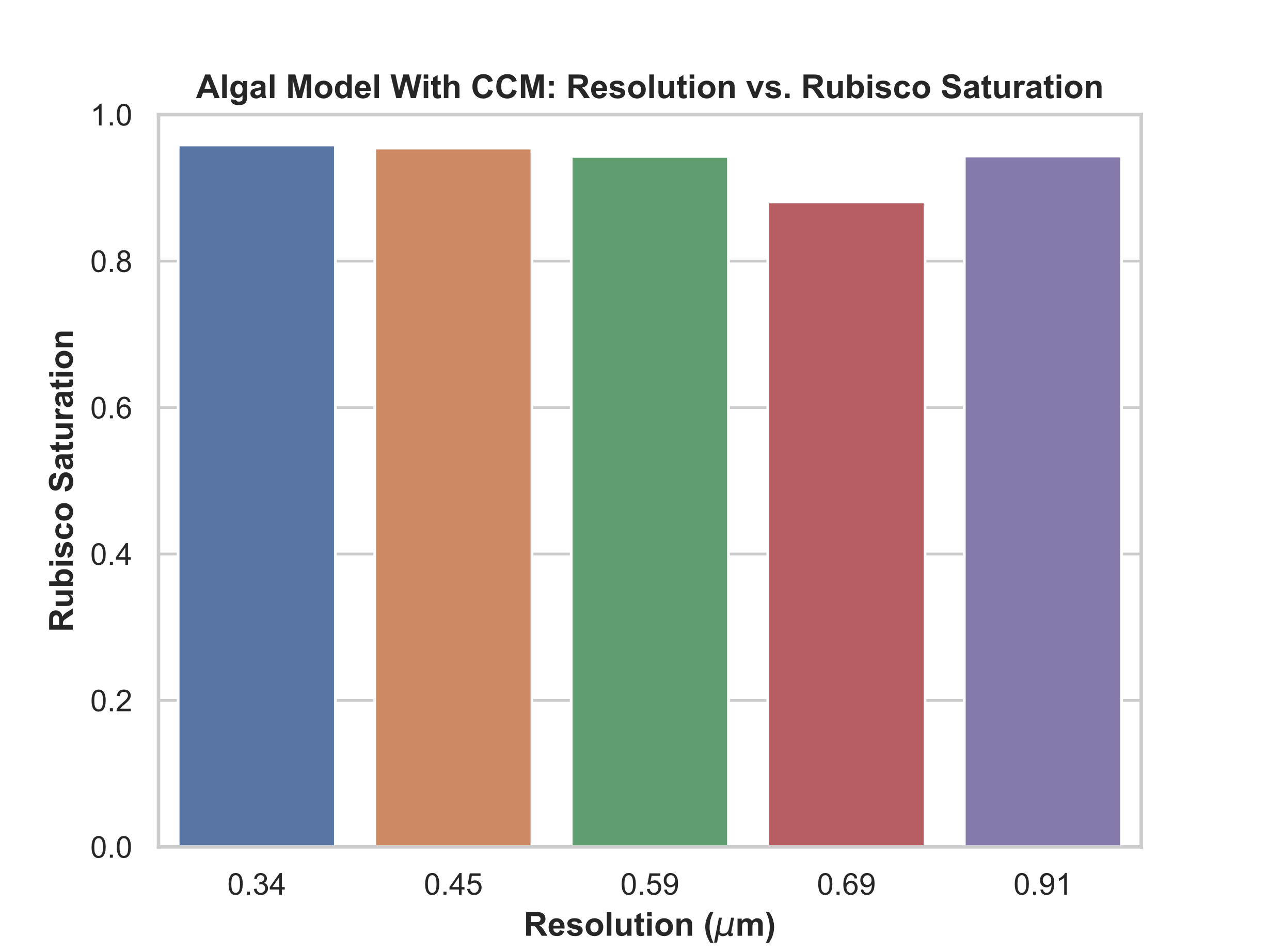

**Figure S4:** Effect of simulation spatial resolution on rubisco saturation. Simulation results are taken from the model of an algal cell with a CCM.

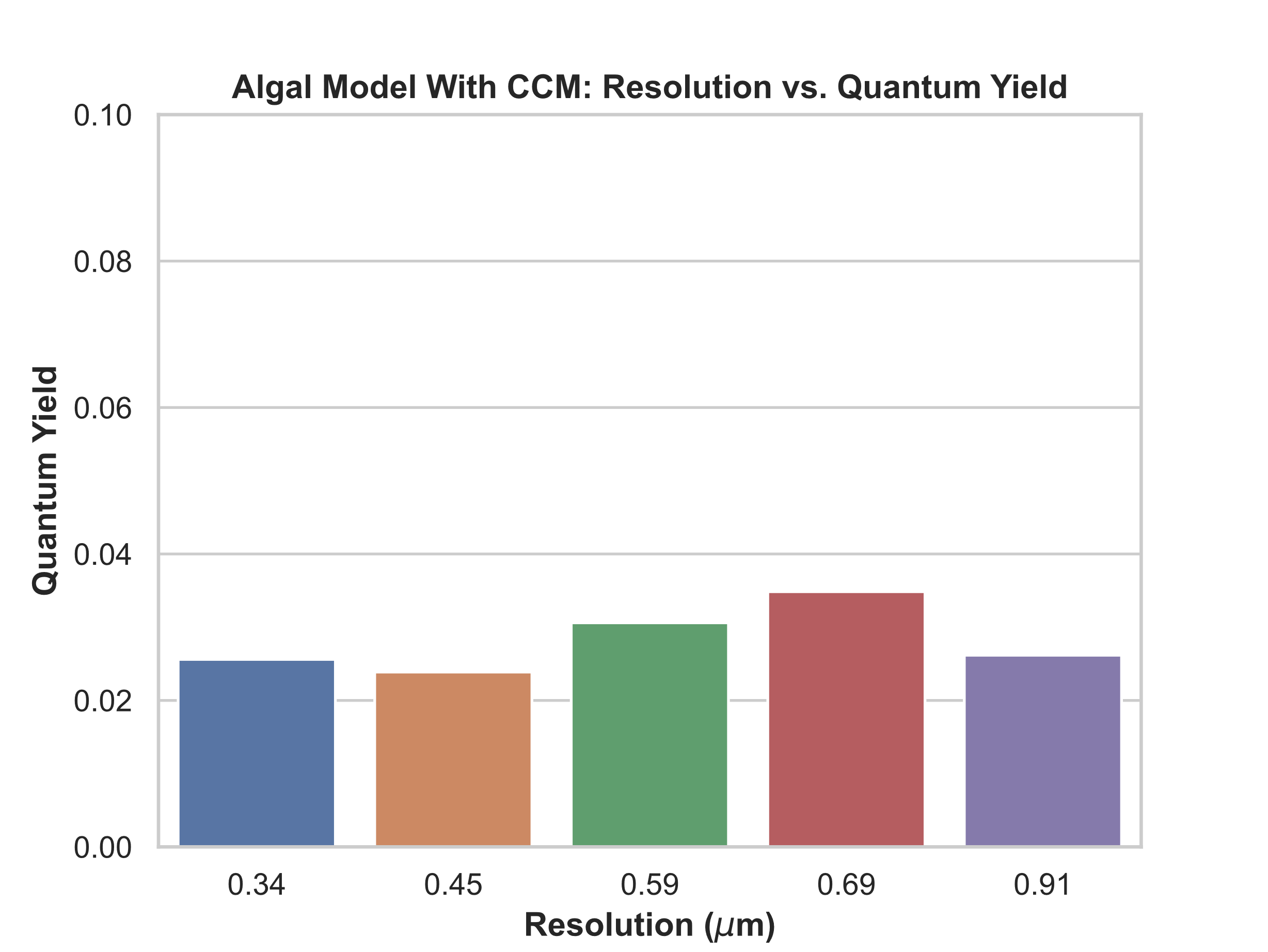
 **Figure S5:** Effect of simulation spatial resolution on quantum yield. Simulation results are taken from the model of an algal cell with a CCM.

**Supplemental References**

1.	B. J. Walker, D. M. Kramer, N. Fisher, X. Fu, Flexibility in the Energy Balancing Network of Photosynthesis Enables Safe Operation under Changing Environmental Conditions. *Plants (Basel, Switzerland)* **9** (2020).
